# VARAN-GIE: Curation of Genomic Interval Sets

**DOI:** 10.1101/333096

**Authors:** Niko Popitsch

## Abstract

Genomic interval sets are fundamental elements of genome annotation and are the output of countless bioinformatics applications. Nevertheless, tool support for the manual curation of these data is currently limited. We developed VARAN-GIE, an extension of the popular Integrative Genomics Viewer (IGV) that adds functionality to edit, annotate and merge genomic interval sets. Data can easily be shared with other users and imported/exported from/to multiple common data formats. VARAN-GIE binary releases, source-code, user guides and tutorials are available at https://github.com/popitsch/varan-gie/

## 1 Introduction

Genomic data sets such as NGS read alignments or SNP array data sets are typically annotated using genomic intervals that define, for example, genes and transcripts, structural and copy number variations or general genomic regions of interest. Such interval sets are usually created by bioinformatics tools such as gene predictors or copy number variation callers but often require subsequent manual curation, e.g., for artefact removal, editing of interval endpoints or merging of adjacent intervals. Curation of genomic interval sets is insufficiently supported by popular genomics browsers such as Broad’s Integrative Genomics Viewer (IGV, Robinson et al. [2011]) or UCSC’s Genome Browser (Kent et al. [2002]) that support mainly read-only access to genomics data and include few basic features for annotating regions of interest therein. Specialized data curation tools, such as ChAS and Nexus Copy Number for copy number variation calling or Apollo Lee et al. [2013] and Artemis Rutherford et al. [2000] for gene curation, support only domain-specific data curation tasks and often lack support for all state-of-the-art file formats or suffer from scalability issues when confronted with huge dataset as produced by whole-genome sequencing (WGS) technologies.

## 2 Approach

Here, we describe VARAN-GIE (VARiant ANnnotation Genomic Interval Editor, aka VARAN, Figure 1) that extends the popular and highly scalable IGV genome viewer with comprehensive functionality for the curation and annotation of genomic interval sets. VARAN enables users to create, modify and merge genomic interval sets that annotate arbitrary genomics datasets loaded into IGV. Comprehensive editing and navigation functionality is directly embedded in IGV’s user interface, e.g., as additional toolbox items and context menus. Basic editing commands include adding, clipping, extending and merging intervals. Commands can be applied to large (sub-) sets of intervals (e.g., all currently visible intervals) and an “undo/redo” mechanism enables users to take back/redo their last actions. Users can create *de novo* interval sets or import and curate data created by upstream bioinformatics pipelines (e.g., BED or tab-separated value/TSV files). Data import can also be done via copy&paste which is convenient when importing from spreadsheets or scientific papers. VARAN uses a “layer” concept for organizing genomic intervals and avoiding visual clutter. Each layer contains a set of genomic segments (herein defined as non-overlapping genomic intervals) and all configured layers are visualized in IGV as BED tracks. Intervals in the currently “active” layer can be curated using the abovementioned tools or copied from any loaded IGV feature track (e.g., gene annotations from a loaded BED file). Intervals can also be moved/copied between different layers and visualized in a special whole-genome view. Curated segment sets from multiple layers can be merged when exporting the data as BED or TSV files which enables authoring of arbitrarily complex (overlapping) interval sets. Genomic intervals are by default associated with meta-data attributes defined in the BED format (name, colour and score) but users can additionally define arbitrary custom meta-data attributes (name/value pairs), e.g., for adding human-readable descriptions or to keep track of numerical values calculated by an upstream bioinformatics pipeline. Interval data can be imported from BED or TSV files (including custom attribute values) after configuring a mapping between TSV headers and interval attributes. Pairs of intervals can be linked to annotate logically connected genomic regions (such as translocation breakpoints or enhancer/TSS regions).Users can then, e.g., quickly navigate between such linked regions in IGV or use this information in downstream processing(e.g., for creating circos plots). Linked intervals can also be imported from BEDPE-formatted files. Taken together, this makes VARAN a powerful tool for the curation and annotation of genomic interval sets.

**Figure 1:**
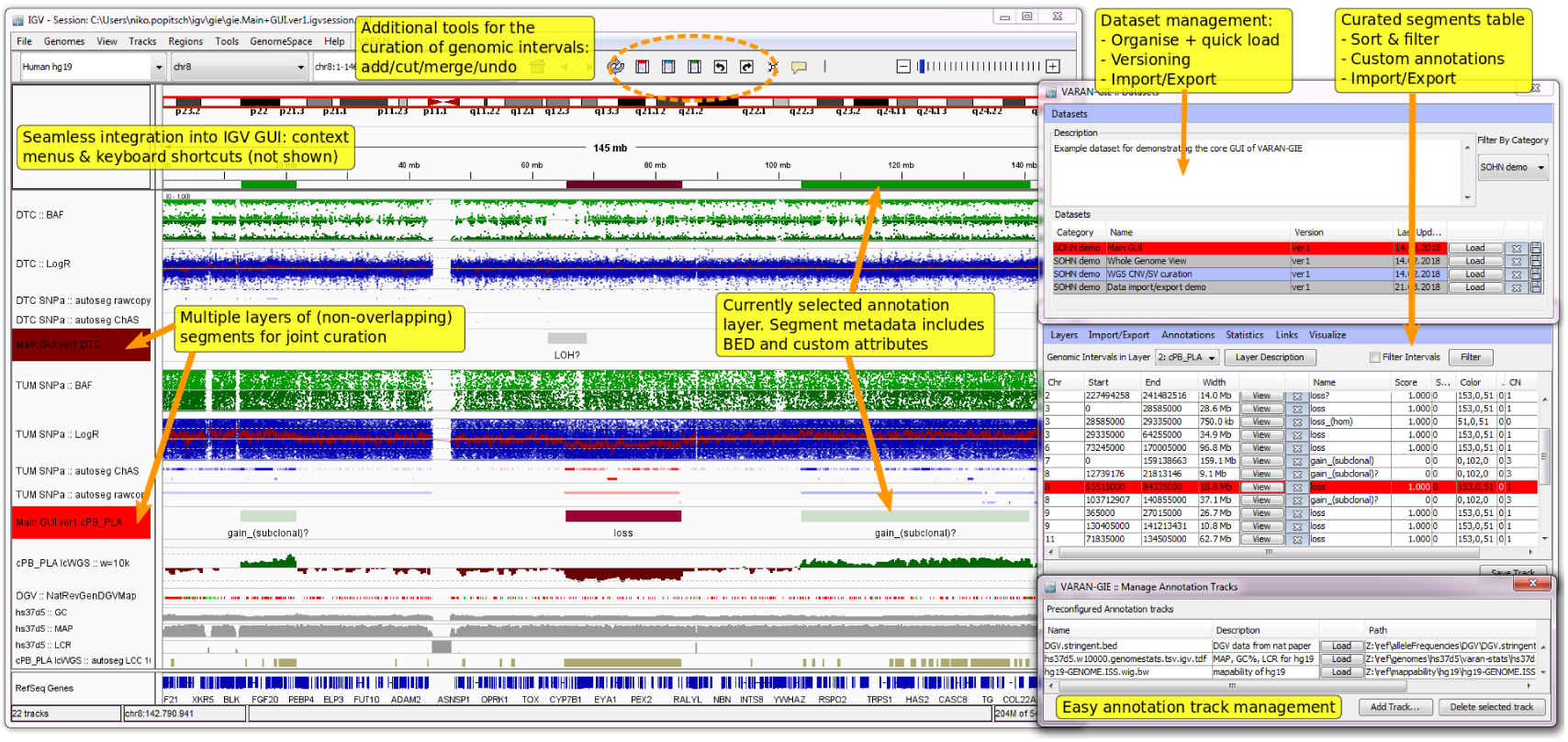
Annotated screenshot explaining the core components of the VARAN GUI and demonstrating how it can be used for the curation of somatic copy-number aberration (SCNA) segments in SNP array and low-coverage WGS data. The window on the left shows the regular IGV GUI that was extended using additional toolbox items and various context menus (not shown). The 3 windows on the right are VARAN-specific and show (from top to bottom) a list of configured datasets, the genomic intervals of the currently selected layer (“cPB PLA”) and a list of configured data tracks than can quickly be loaded by the user. The shown IGV data tracks contain SNP array and low-coverage WGS data as well as several annotation tracks. The two highlighted BED tracks are automatically maintained by VARAN and contain intervals configured in two different data layers. The currently active layer is highlighted in bright red color and the respective intervals are also shown in the top panel just below the chromosome ideogram. Clicking on these intervals opens context menus for interval curation (not shown).

Intervals of the currently active layer can also be edited in a filterable and sortable data table. This enables users to select interval subsets and apply common changes such as setting the values of a particular meta-data attribute. The data table can be filtered by genomic scope (e.g., to show only intervals currently visible in IGV) and by sequentially applied, attribute-based filter expressions (e.g., to display only autosomal intervals longer than a particular threshold and above a given score). Filtered intervals can then be exported to 3rd party applications via copy & paste. Whole interval layers can be exported as BED or TSV files, whole datasets (including all layers) as small ZIP files that contain interval sets, configuration data and an IGV session file that contains references to the respective genomic data files. Such ZIP files can easily be shared with colleagues (e.g., via email) which can then import and view the data exactly as it was exported given they have access to the respective genomics data. Broken file references can be fixed at import time via a special dialog, see online documentation for details. Overall, this functionality effectively enables the collaborative curation of interval sets that annotate arbitrary genomics data.

VARAN supports dataset versioning and organization in arbitrary categories and users can quickly (“one-click”) switch between datasets. Users are protected from data loss by an automatic autosave functionality that enables them to simply restart the application at any time and continue their work where they left it in case IGV became unresponsive or crashed. VARAN was developed by branching IGV version 2.3.94 and extending the Java source-code with application-specific code.

## 3 Conclusion

VARAN is a universal and highly scalable approach for the curation of genomic intervals. Use cases include curation of structural variation segments in WGS data, *de-novo* annotation of transcripts in RNA-seq data or merging and post-processing of interval sets resulting from bioinformatics pipelines. We are currently using VARAN for the integrative curation and comparison of somatic copy-number aberrations in spatially and temporally diverse neuroblastoma patient samples to obtain deeper insights into intra-tumour heterogeneity, especially when comparing tumour DNA with cfDNA interrogated on different platforms (e.g., SNP array, low-coverage and deep WGS).

## Acknowledgements

The author would like to thank Teresa Gerber and other members of the Ambros group at CCRI for providing data and stringently testing VARAN.

## References

J. W. J. Kent, C. W. Sugnet, T. S. Furey, K. M. Roskin, T. H. Pringle, A. M. Zahler, and D. Haussler. The human genome browser at ucsc. Genome research, 12(6):996–1006, 2002.

E. Lee, G. A. Helt, J. T. Reese, M. C. Munoz-Torres, C. P. Childers, R. M. Buels, L. Stein, I. H. Holmes, C. G. Elsik, and S. E. Lewis. Web apollo: a web-based genomic annotation editing platform. Genome biology, 14(8):R93, 2013.

T. Robinson, H. Thorvaldsdóttir, W. Winckler, M. Guttman, E. S. Lander, G. Getz, and J. P. Mesirov. Integrative genomics viewer. Nature biotechnology, 29(1):24, 2011.

Rutherford, J. Parkhill, J. Crook, T. Horsnell, P. Rice, M.-A. Rajandream, and B. Barrell. Artemis: sequence visualization and annotation. Bioinformatics, 16(10):944–945, 2000.

